# OVEREXPRESSION OF TUMORPROTECTIVE CLUSTERIN PREVENTS EXHAUSTION OF TRANSGENIC T CELLS

**DOI:** 10.64898/2025.12.16.694090

**Authors:** Constantin Segner, Yue Z Huang, Elena Sotillo, Alisa Kolesnikova, Mansour Poorebrahim, Julia Höbart, Michael Lizardo, Julia Mergner, Fares Burwag, Miriam Schulz, Busheng Xue, Andrew Minchinton, Christian Brückner, Alissia Fernandes Madeira, Julia Hauer, Crystal Mackall, Jürgen Ruland, Poul H. Sorensen, Stefan E.G. Burdach

## Abstract

T cell therapies, such as chimeric antigen receptor (CAR) T cells and T cell receptor (TCR) transgenic (tg) T cells, are a promising approach in the treatment of solid malignancies but are limited by T cell exhaustion caused by chronic antigen stimulation. Clusterin (CLU) is a chaperone protein known to protect both normal and malignant cells, from metabolic stress and reactive oxygen species (ROS). In this study, we investigated whether overexpression (OE) of CLU in TCRtg T cells and CAR-T cells respectively can reduce exhaustion induced by chronic antigen stimulation and enhance T cell functionality against Ewing sarcoma (EwS). Among other cytoprotective genes, we found that CLU was significantly downregulated in dysfunctional tumor infiltrating lymphocytes. Therefore, we engineered EwS-directed TCRtg T cells targeting a Chondromodulin-1 (Chm1)-derived peptide and GD2 CAR-T cells to overexpress CLU. We show here that CLU is downregulated in T cells following activation by tumor cells. CLU-overexpressing T cells exhibit decreased expression of exhaustion markers (PD1, LAG3) and reduced apoptosis after repetitive stimulation. These cells demonstrated improved infiltration into tumor spheroids and maintained functionality under hypoxic conditions. *In vivo*, CLU-overexpressing T cells showed enhanced persistence and a trend towards reduced tumor growth. Mechanistically, proteomic analysis suggested that reduced ribosomal activity might delay T cell exhaustion, implicating metabolic reprogramming.

In conclusion, CLU OE in tg T cells enhances their persistence and functionality by mitigating exhaustion, possibly through modulation of ribosomal activity and metabolic pathways. This strategy holds potential for improving adoptive T cell therapies against solid tumors.

## 3. Introduction

Immunotherapy has been a recent breakthrough in cancer treatment and T cells are main players in this scenario. Antibody-based strategies to armor T cells against cancer involved bypassing physiological immune breaks, which naturally limit the intensity and duration of T cell attacks (e.g. anti-CTLA4 antibodies, e.g. Ipilimumab or anti-PD1 antibodies, e.g. Nivolumab).^1, 2^ In addition, cell-based strategies of targeting tumor cells, by circumventing T cell restriction, has been achieved by short circuiting the evolutionary dichotomy of acquired immunity into its humoral and cellular branches. The latter has been accomplished by inserting immunoglobulin genes fused to genes encoding the T cell receptor signal transduction and costimulatory machinery into the T cell genome to the effect, that the T cell serves as a payload for the immunoglobulin molecule which recognizes the target structures on the tumor cell membrane.^3^ These chimeric antigen receptor (CAR) T cells - as well as transgenic (tg) TCR T cells - provide an exciting new avenue in the development of immunotherapies.^4^

One major limitation of T cell therapies in cancer treatment is the differentiation of T cells into an exhausted phenotype (T_ex_) upon constant activation tonic signaling of the T cell receptor.^5^.^6–8^ Besides that, a hyperactive Akt-mTOR-S6K pathway and increased protein synthesis promote early T cell exhaustion.^9^ T_ex_ exhibit a significantly impaired effector functionality.^10^ These functional changes are associated with expression of surface markers can be used to identify exhausted T cells.^5^ (e.g. PD1^11^, LAG3^12^, TIM-3, 2B4 and TIGIT^13^).

Clusterin (CLU) is a chaperone, which has been recognized to ameliorate stress in both intracellular and extracellular environments.^14^ CLU’s role lies in stabilizing the post-translational structure of various proteins and preventing the aggregation of misfolded proteins.^15, 16^ Thereby, it protects cells, e.g., in the context of tumors, from metabolic stress and excessive production of reactive oxygen species (ROS).^17^ It is known that the metabolic state in T cells plays an essential role in their transition to an exhausted state.^18, 19^

Ewing sarcoma (EwS) is a highly malignant tumour of the connective tissue, which affects mostly adolescents in early puberty and tends to metastasize early. In advanced stages, the disease has a dismal prognosis, which funds an urgent need for efficacious therapies.^20, 21^ In addition, because of its low mutational burden critical target lesions are easier to identify in EwS as compared to the common high mutational burden cancer of the Old. Thus, albeit a rare disease, it may serve as an excellent model for metastasizing cancer, confirming the paradigm that rare diseases may pave the road to precision medicine.^22^

Recently, we have provided preliminary safety and efficacy data of immunotherapy of patients with advanced EwS utilizing tgTCR-T cells targeting HLA restricted peptides derived from metastatic drivers.^23^ In preceding preclinical work, we identified Chondromodulin-1, encoded by the LECT1 gene, as being uniquely upregulated in EwS, compared to the normal body expression atlas, using microarray analysis.^24, 25^ Based on our further characterization of Chm1 in EwS, in particular its role in metastatic spread,^26^ as well as the identification of an EwS selective Chm1-derived immunogenic peptide and the cloning of its cognate TCR, we revealed potential of Chm1 as a suitable target for specific TCR based therapy.^25 27, 28^. In addition to Chm1, the disiaganglioside GD2 has emerged as another promising target for immunotherapy in EwS. Recent studies have shown that GD2 is expressed in 40% to 90% of EwS cells, making it a suitable tumor-associated antigen for CAR-T cell therapies.^29, 30^

In this study we tested the hypothesis that an ectopic OE of CLU in multiple adoptive T cell (e.g. Chm1 targeting tg T cells) therapy model, can ameliorate the detrimental effects of chronic antigenic stimulation and improve the T cells functionality within the TME.

## 4 Materials and Methods

### 4.1 Cell lines, reagents, and media

The cell lines utilized in this study have been described previously.^31, 32^ The IL15-producing myeloma line NSO was a kind gift from Prof. Stanley Riddell. Ewing sarcoma lines A673, TC71 and GD2/luciferase-expressing Nalm6 (gift from Prof. Rob Majzner) were authenticated and mycoplasma-tested. Cells were cultured in RPMI 1640 with 10% FBS and penicillin/streptomycin. Packaging lines 293Vec-RD114 and HEK293 (gift from Prof. Manuel Caruso) were maintained in DMEM with 10% FBS, penicillin/streptomycin, sodium pyruvate, and non-essential amino acids. Human T cells were expanded in T Cell Expansion Medium (StemCell Technologies) supplemented with 5% human serum, 200⍰mM N-Acetyl-L-Cysteine, penicillin/streptomycin and IL-2 (10⍰IU/mL). PBMCs were isolated from healthy DRK donors (18–65⍰years; screened for HIV, HBV, HCV, syphilis) by Ficoll-Paque density-gradient centrifugation, following Ethics Committee approval (protocol #XYZ) and written informed consent.

### 4.2 Expansion of GD2 CAR-T cells and tgTCR-T Cells

To generate antigen-specific tg T cells, ChoTCR and ChoTCR^CLU^ T cells were produced using a retroviral vector encoding the murinized Chondromodulin-1 TCR, with the Clusterin sequence kindly provided by Prof. Poul Sorensen. GD2 CAR-T cells were created with a lentiviral vector incorporating GFP as a transduction marker. After a week of expansion, ChoTCR and ChoTCR^CLU^-T cells were purified by staining with a PE-conjugated antibody specific to the murinized TCR, while GD2 CAR-T cells were sorted for GFP expression. Cell sorting was performed on a BD FACSAria Fusion flow cytometer (BD Biosciences, Heidelberg, Germany). To facilitate the expression of the transgene, we used a pMP71 plasmid encoding for the MP71 retroviral backbone and the ChoTCR. We fused the T2A_CLU construct to the 3’ end of the TCR using the In-Fusion® Snap Assembly Kit from Takara according to the manufacturer’s instructions **(Supplementary Figure 1A-C)**. The generation of the CLU.GD2 plasmids was achieved in a similar manner. Cell lines, media and reagents are listed in **Supplementary Table S1**.

Purified ChoTCR and ChoTCR^CLU^ T cells were cultured in 25 cm^2^ flasks (TPP, Trasasingen, Switzerland) with 25 mL T cell medium supplemented with OKT3 (50 ng/mL; Biolegend, San Diego, CA, USA), IL-2 (100 IU/mL; R&D Systems, Minneapolis, MN, USA), IL-7 (5 ng/mL; R&D Systems), and IL-15 (2 ng/mL; R&D Systems), refreshing interleukins every week.

### 4.3 Flow cytometry-based cell characterization

For FACS analysis, cells were harvested at equal numbers.Surface antigen staining was performed with a 1:300 dilution of the primary antibody in FACS buffer (1% BSA in PBS) for 30 minutes at 4°C, followed by secondary antibody staining when necessary. For intracellular antigen analysis, cells were fixed in 4% formaldehyde, permeabilized with ice-cold 90% methanol, and stained as above. Analyzes were conducted on a BD FACSCanto II or BD LSRFortessa, with compensation managed in FlowJo (RRID:SCR_008520). All antibodies are listed in **Supplementary Table S2**.

### 4.4 Analysis of Cytokine Production Using Enzyme-Linked Immunosorbent Assay (ELISA)

To analyze IL-2 and IFN-γ production in GD2 CAR T cells, enzyme-linked immunosorbent assays (ELISAs) were performed using BioLegend ELISA MAX^TN^ Standard Sets according to the manufacturer’s instructions.

### 4.5 Lactate Dehydrogenase (LDH) Release Assay to determine cytotoxicity

To assess the cytotoxicity of Chondromodulin-1 targeting T cells, we used the Lactate Dehydrogenase (LDH) Activity Assay Kit from Sigma-Aldrich according to the manufacturer’s instructions. This assay quantifies LDH release as a marker of cell damage and cytotoxicity.

### 4.6 Bioluminescence Assay to determine cytotoxicity of GD2-CAR T cells

To assess GD2 CAR-T cell cytotoxicity, we used the Nano-Glo® Luciferase Assay System (Promega) with luciferase- and GD2-expressing Nalm6 target cells. Nalm6 cells were co-cultured with GD2 CAR-T cells for varying durations, depending on the specific experimental conditions. Supernatants were analyzed according to the manufacturer’s instructions.

### 4.7 Confocal Microscopy

Confocal microscopy was performed on a Zeiss LSM 800 Confocal Laser Scanning Microscope. T cells were stained with either PE or Alexa Fluor 647 conjugated anti-CD3 antibodies (RRID:AB_314043 and RRID:AB_571883) before starting the co-culture at a concentration of 1:15. EwS cells with transgenic expression of GFP were used to visualize the spheroids. Images were captured at a magnification of 20x. For acquisition of z-stacks, the lower apex of the spheroid was acquired, then 25 to 35 images towards the upper apex of the spheroid were taken. 3-dimensional z-stack analysis was performed using the Imaris Microscopy Image Analysis Software from Oxford Instruments (RRID:SCR_007370).

### 4.8 Animal Experiments

All animal experiments were approved by the University of British Columbia Animal Care Committee (protocol# A23-0167) and conducted following institutional ethical guidelines. Rag2-/-γc-/-mice (BALB/c background; RRID:IMSR_JAX:014593; Stock No: 014593) were obtained from The Jackson Laboratory (Bar Harbor, Maine, USA) bred at the British Columbia Cancer Research Center under pathogen-free conditions, with experiments performed on 12-week-old female mice. A673-GFP-Luc EwS cells (5×10^5^) were injected intramuscularly (i.m.) into the right calf, and mice were randomized into treatment (ChoTCR or ChoTCRCLU; n=8) or control (NT; n=4) groups. No blinding was performed. Mice were screened prior to enrollment to ensure they were healthy and free of visible abnormalities. No mice met pre-defined exclusion criteria (infection or injury), and all enrolled animals proceeded to the experimental phase. Eight days post-tumor cell injection, mice received 5×10^6^ T cells (NT, ChoTCR, or ChoTCRCLU) derived from healthy human donor PBMCs, administered intraperitoneally (i.p.). All mice were injected i.p. biweekly with 1×10^7^ irradiated (100 Gy) IL15-NSO cells. Mice were monitored biweekly for weight, tumor size, and general health. Humane endpoints were defined by a scoring system that included tumor size (>17 mm diameter), ulceration, significant weight loss, respiratory distress, reduced activity, or behavioral changes. Ten days post-infusion, one NT and two treated mice were sacrificed for phenotype analysis; the rest were euthanized at endpoints. Tumor size was assessed weekly through bioluminescence; mice were injected i.p. with 150 mg/kg luciferin, and total photon flux was measured 10 minutes later on an IVIS Lumina (Caliper Life Sciences, Waltham, MA) using automated exposure detection. Mice were briefly anesthetized with 1.5% isoflurane for imaging. At endpoints, mice were euthanized by isoflurane anesthesia, CO_2_ inhalation, and cervical dislocation. No blinding was performed. Explanted organs were fixed in 4% formaldehyde. Spleens were mechanically dissociated, treated with RBC Lysis Buffer, and filtered for flow cytometry. Other tissues were fixed in formaldehyde for up to 96 hours before transfer to 70% ethanol for long-term storage. CD99 and CD3 staining in lung tissues were analyzed using QuPath (version 0.5.1). The cell detection tool enabled automated cell identification, with a higher CD99 threshold set to distinguish Ewing sarcoma cells from T cells.^33^ The required sample size for the Mann-Whitney U test, assuming a power of 0.8, was calculated to be n = 46 per group. Therefore, a Bayesian t-test was chosen as it accommodates smaller sample sizes and provides robust statistical inference.

### 4.9 Proteomic Analysis

Mass-spectrometry-based proteomics analysis was conducted at the Bavarian Center for Biomolecular Mass Spectrometry at Klinikum rechts der Isar (BayBioMS@MRI). Cell pellets were lysed in urea extraction buffer (8M Urea, 50 mM Tris-HCl, pH 7.5), followed by sonication and centrifugation. Protein concentration was determined via Bradford assay, and samples were digested with trypsin after reduction with dithiothreitol and alkylation with chloroacetamide. Peptides were labelled with TMT10plex reagent as described previously.^34^ A Dionex Ultra 3000 HPLC system operating a Waters XBridge BEH130 C18 3.5⍰µm 2.1⍰×⍰250⍰mm column was used to fractionate the TMT sample. 96 fractions were collected, pooled to 24 fractions, acidified to 1% with formic acid and dried down prior to LC-MS/MS analysis. Nanoflow LC-MS/MS was performed on a Dionex 3000 (Thermo) coupled online to an Orbitrap Eclipse (Thermo) mass spectrometer. Samples were measured in DDA mode with a MS3 method and 50 min linear gradient. Database search against the human reference proteome (UP000005640) and common contaminants was performed with MaxQuant (v. 2.3.0.0) with standard settings for reporter ion MS3.^35^

### 4.10 Bulk RNAseq Data Analysis

All statistical analyses and graphical visualizations were conducted in R (version 4.2.0). Data were imported using the readxl package (Wickham & Miller, 2022). For data manipulation and transformation, we used the dplyr (Wickham et al., 2022) and tidyr (Wickham, 2022) packages. All plots, including Volcano and Bar Charts with error bars and significance annotations, were generated with ggplot2 (Wickham, 2016; RRID:SCR_014601) and further refined using ggrepel (Slowikowski, 2020) to improve label readability.^36–39^

### 4.11 Single-Cell RNA-Sequencing Analysis

Single-cell RNA-seq data were processed using the Seurat R (RRID:SCR_007322) package. Briefly, cells with low gene counts or high mitochondrial content were filtered out. Data were normalized using a log-transformation and scaled prior to dimensionality reduction. Principal component analysis (PCA) was performed, and the top principal components were used to generate UMAP embeddings for visualization.

T-cell subpopulations were defined using according to the Immune Cell signatures published by Monaco et al.^40^ All Visualizations were created using R as described above.

### 4.12 Statistical Analysis and Visualization

All charts were generated using FlowJo, GraphPad Prism (RRID:SCR_002798), OriginLab (RRID:SCR_014212) or R as described above.^41–43^

## 5 Results

### 5.1 Cytoprotective Proteins, Including CLU, are Downregulated in Exhausted Tumor Infiltrating Lymphocytes

To identify cytoprotective genes potentially involved in safeguarding T cells from exhaustion, we reanalyzed publicly available scRNAseq expression data from Sade-Feldman et al., who characterized transcriptional profiles of tumor-infiltrating CD8^+^ T cells from melanoma patients receiving checkpoint blockade therapy. Building on these findings, we utilized the Immune Reference from Monaco et al.^40^ to delineate immune cell subpopulations and evaluated the expression of 12 cytoprotective genes known to confer resistance to metabolic and therapy-induced stress in solid tumors **(Figure 1A)**.

**Figure 1.**
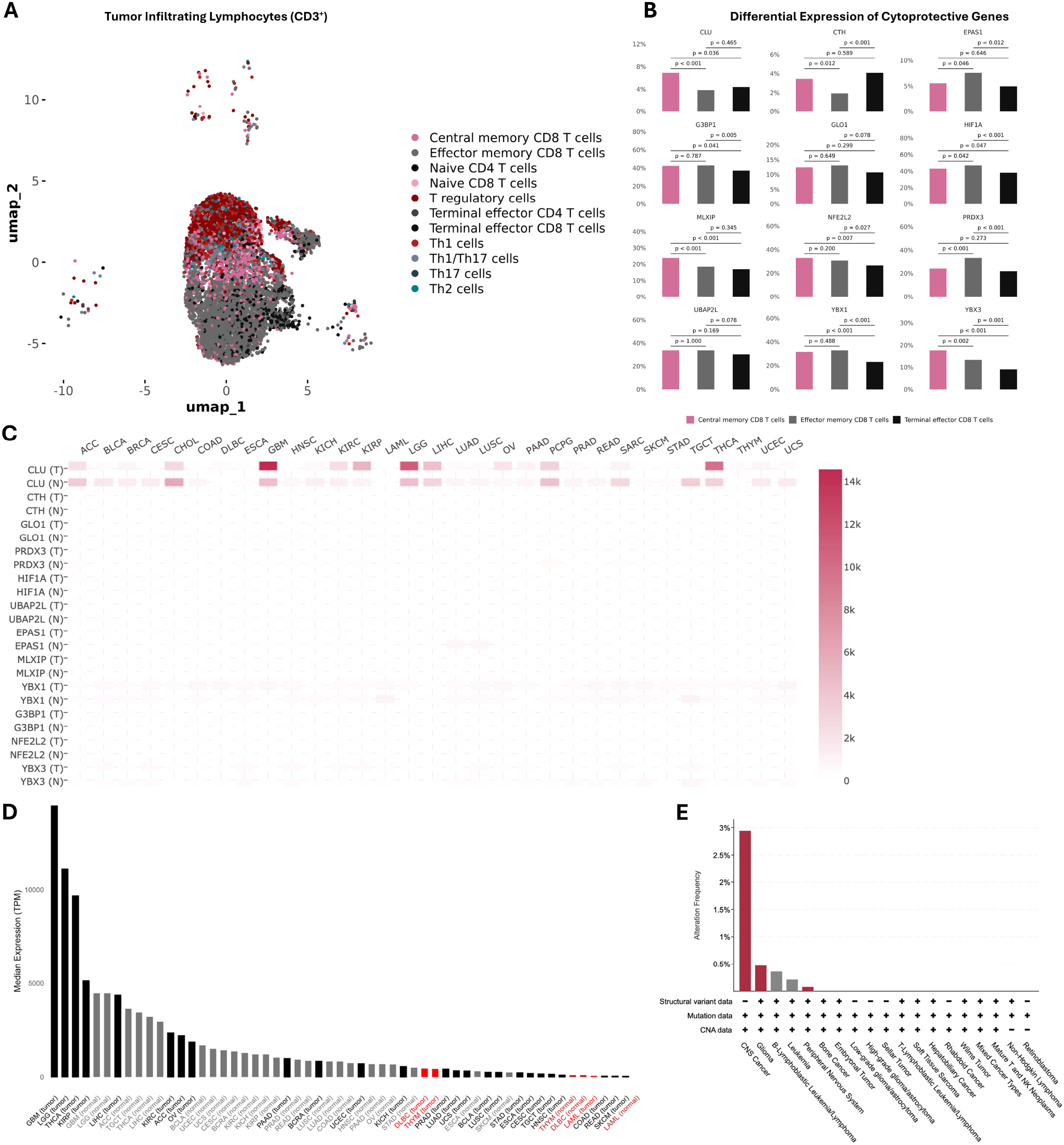
Cytoprotective Protein Screening in Tumor Infiltrating Lymphocytes. (**A**) UMAP visualization of five T-cell subpopulations (Naive, Central Memory, Effector Memory, Cytotoxic, and Exhausted) reanalyzed from GSE120575. The analyzed data represents Tumor-Infiltrating Lymphocytes from Melanoma patients, after receiving immune-checkpoint inhibitor treatment (**B**) Combined violin plots illustrating the expression of 12 cytoprotective genes in five T-cell subpopulations (Wilcoxon test p-values annotated for Naive versus Exhausted) using the same color scheme as in (A), reanalyzed from GSE120575. **(C)** Expression of cytoprotective genes in cancer (T) and healthy (N) tissue. TPMs are presented in linear scale. Data extracted from cbioportal. (**D**) CLU expression in different healthy and malignant human tissues (“normal” refers to the corresponding tissue of origin) Hematopoietic tissues are displayed in red. Data sourced from GEPIA. (**E**) Percent pediatric patients with mutations in the CLU gene in all tumors. Red denotes mutation, grey denotes deep deletion. Data extracted from cbioportal.org

Out of these analysed genes CLU, G3BP1, HIF1A, MLXIP, NFE2L2, YBX1 and YBX3 were downregulated in Terminal Effector CD8^+^ T cells compared to Central Memory CD8^+^ T cells, while CLU, MLXIP and YBX3 were also downregulated when comparing Effector Memory CD8^+^ T cells with Central Memory CD8^+^ T cells **(Figure 1B)**. Next, we looked at the expression levels of our cytoprotective gene set in healthy and tumor tissue (GEPIA) and observed that CLU is the highest expressed of the 12 cytoprotective genes probed both in cancer and normal tissue **(Figure 1C)**. We hypothesize that CLU overexpression (OE) in T cells would have the strongest advantageous phenotypic effect, while retaining a non-oncogenic profile. CLU is a chaperone protein with essential functions in lipid transport, metabolism, and stress response. It regulates metabolic homeostasis, insulin sensitivity, and adipocyte function, while also protecting tumors from metabolic stress.^17, 44, 45^ To assess the potential risk of a Clusterin induced transformation in a CAR-T cell, we assessed the expression of CLU using GEPIA in human tissue, both normal and malignant. CLU was highly upregulated in certain solid tumors (e.g., glioblastoma multiforme and thyroid carcinoma), with expression levels far exceeding those observed in CAR-T cell models. On the other hand, the expression of CLU in hematopoietic tissues and tumors was very low **(Figure 1D)**. To further address the role of CLU in progression of pediatric malignancies, we screened the TCGA dataset for structural changes in the CLU gene (cBioportal.org). In pediatric cancer we found no amplifications and only few deep deletions and mutations insinuating that even if CLU is being upregulated, it rather acts as a passenger than a driver mutation in these instances **(Figure 1E)**. These findings collectively suggest that CLU OE in T cells can mitigate exhaustion, reduce apoptosis, and enhance cytotoxicity, making it a promising safe strategy to improve the efficacy of T cell-based immunotherapies.

### 5.2 CLU OE Rescues T Cell Functionality After Prolonged Stimulation

Previously, our group developed an HLA-A^*^02:01/CHM1-specific allorestricted T cell receptor (ChoTCR). We overexpressed CLU using a polycistronic vector in ChoTCR-T cells **(Figure 2A)**. Retroviral transduction with the CLU gene led to a significantly increased expression of the CLU protein, as measured by intracellular flow cytometry **(Figure 2B)**. Next, we investigated the impact of CLU OE on the functional and cytotoxic properties of ChoTCR-T cells after 24 hours of stimulation with tumor cells. Although ChoTCR-T cells with CLU OE exhibited lower levels of IFN-γ, a key cytokine involved in anti-tumor responses, there was no significant difference in Granzyme B (GzmB) production and anti-tumor killing activity **(Figure 2C-D)**. To evaluate the effect of CLU-OE on the expression of inhibitory receptors upon stimulation, we challenged non-transduced (NT), ChoTCR, and ChoTCR^CLU^-T cells with chondromodulin-I/HLA-A^*^02:01 expressing EwS cell lines A673 and TC71. We observed a significantly decreased expression of the inhibitory receptors PD-1 and LAG3 in ChoTCR^CLU^-T cells upon stimulation with both cell lines and at different timepoints **(Figure 2E-F)**. Furthermore, the ChoTCR^CLU^-T cells showed reduced expression of the apoptosis marker Annexin V, improved persistence in co-culture and increased Ki67 expression after stimulation with A673 EwS cells for 6 days **(Figure 2G and Supplementary Figure 2A-B)**. This reduction in apoptosis indicates that CLU OE may confer a survival advantage to T cells upon activation by tumor cells. T cell exhaustion progresses hierarchically through four stages—quiescent T_ex_ progenitor 1 (SLAMF6^+^CD69^+^), cycling T_ex_ progenitor 2 (SLAMF6^+^CD69^−^), effector-like intermediate (SLAMF6^−^CD69^−^), and proliferation-incapable terminal (SLAMF6^−^CD69^+^)—with TOX expression peaking in the terminally exhausted subset. Next, we stained with SLAMF6 (Ly108) and CD69 to determine the degree of exhaustion **(Figure 2H)** and observed that 52.4% of ChoTCR^CLU^-T cells remained in the progenitor 1 and 2 stages following long-term stimulation with tumor cells. In comparison, only 21.3% of ChoTCR-T cells exhibited these phenotypes under the same co-culture conditions. More importantly, only 3.4% of ChoTCR^CLU^-T cells exhibited the Tex terminal phenotype, compared to a substantially higher 30.1% in ChoTCR-T cells. ChoTCR-T and ChoTCR^CLU^-T cells showed a comparable percentage of Tex intermediate phenotype **(Figure 2I)**. These data suggest that CLU OE helps maintain T cells in a less exhausted state, preserving their proliferative and functional potential. Finally, we assessed the cytotoxic potential in a rechallenge assay, whereby T cells pre-stimulated with A673 EwS cells for 9 days were co-cultured a second time. After 24-hours, it was evident that the prestimulated ChoTCR^CLU^-T cells had a better cytotoxicity profile than ChoTCR T cells upon restimulation, which can be attributed to their less exhausted phenotype **(Figure 2J)**.

**Figure 2.**
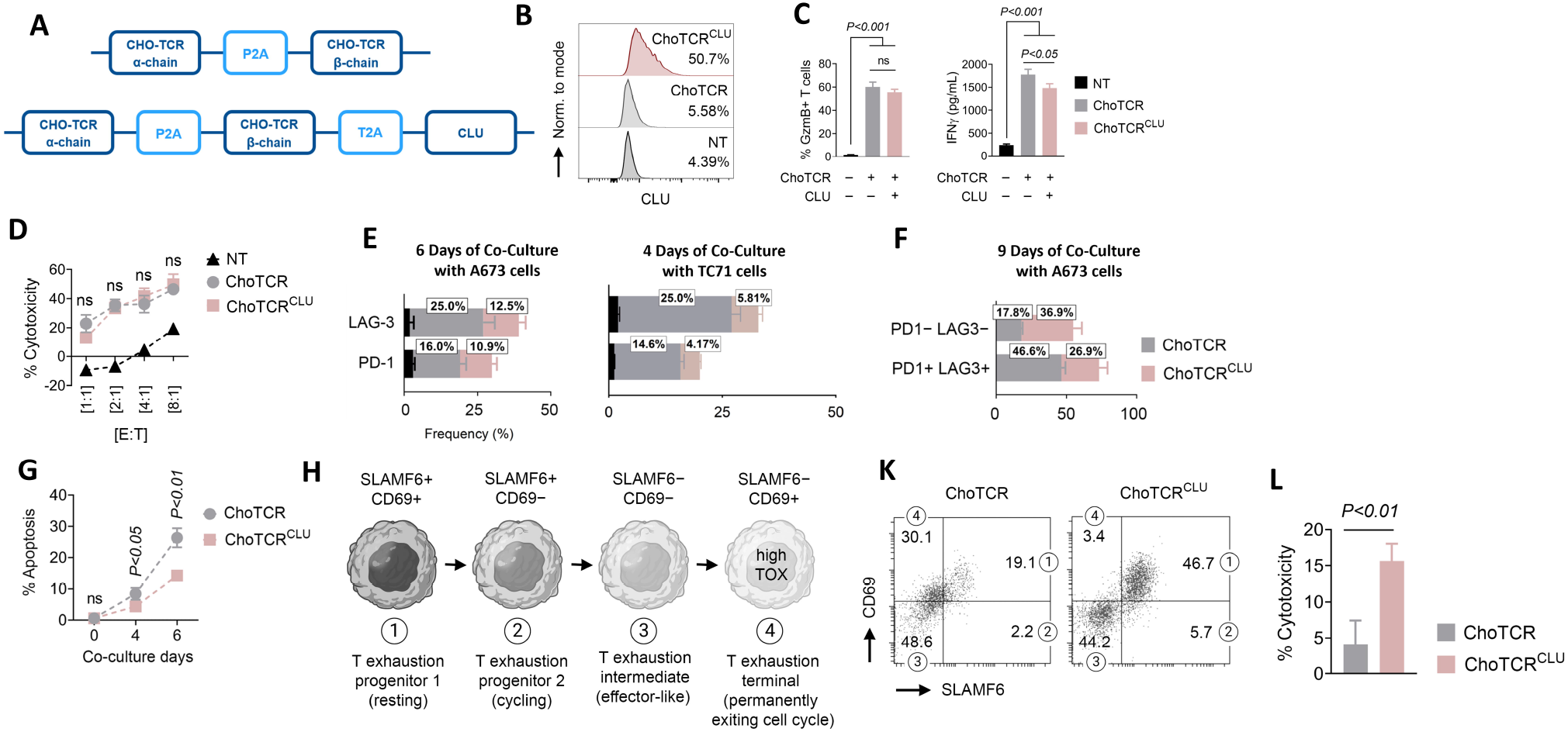
CLU Delays the Transition of tgTCR T cells into an Exhausted State. (**A**) Schematic representation of the pMP71 plasmid coding for the ChoTCR and ChoTCR^CLU^ T cells. (**B**) CLU expression analysis in non-transduced (NT), ChoTCR-T, and ChoTCR^CLU^-T cells. (**C**) Analysis of GzmB and IFN-_γ_ production in the NT, ChoTCR-T, and ChoTCR^CLU^-T cells following 24h co-culture with A673 tumor cells (E:T=1:2). ns: non-significant (P>0.05) (**D**) Lactate dehydrogenase (LDH) cytotoxicity assessment of NT, ChoTCR-T, and ChoTCRCLU-T cells after 24h co-culture with A673 tumor cells at different E:T ratios. ns: non-significant p-values between ChoTCR-T and ChoTCRCLU-T groups. (**E-F**) Analysis of expression of PD1 and LAG3 in the ChoTCR-T and ChoTCRCLU-T cells after stimulation with A673 and TC71 tumor cells (E:T=1:10). Percentages show the average values in ChoTCR-T and ChoTCR^CLU^-T cells. (**G**) Percentage of apoptosis as calculated from ChoTCR+ Ann.V+ populations in the ChoTCR-T and ChoTCR^CLU^-T cells following stimulation with A673 tumor cells. T cells were first gated based on the ChoTCR expression in all comparative flow cytometry analyzes. (**H**) Schematic representation of T cell exhaustion transitions based on the expression of SLAMF6 and CD69. The highest level of TOX (an exhaustion transcription factor) is observed in the terminally exhausted subsets. (**I**) Phenotypic analysis of ChoTCR-T and ChoTCR^CLU^-T cells after third stimulation by A673 tumor cells. The numbers in the circles refer to the transition stage of T cell exhaustion as described in part “C”. The numbers inside the plot represent the percentage of each population. (**J**) Lactate dehydrogenase (LDH) cytotoxicity assessment of ChoTCR-T and ChoTCR^CLU^-T cells after a subsequent stimulation after 9 days, again with A673 tumor cells (E:T=1:2).

### 5.3 CLU OE Improves Tumor Spheroid Infiltration

To investigate ChoTCR-T and ChoTCR^CLU^-T cells cytotoxic function in 3D tumor models, we established tumor spheroids using GFP-expressing A673 EwS cells. These 3D models mimic the *in vivo* tumor microenvironment more closely than 2D cell cultures. Engineered or non-targeting (NT) T cells were added 24h after cell seeding, allowing establishment of structured tumor masses before encounter with T cells. After 24h of co-culture, both ChoTCR-T and ChoTCR^CLU^-T cells induced spheroid destruction, whereas spheroids co-cultured with NT cells preserved their shape and integrity **(Figure 3A)**. Calculation of corrected total cell fluorescence of all analyzed spheroids in co-cultures containing NT (n=14), ChoTCR-T (n=19), and ChoTCR^CLU^-T cells (n=18) demonstrated no significant differences between the killing activity of ChoTCR-T and ChoTCR^CLU^-T cells **(Figure 3B)**. This quantitative analysis suggests that the overall cytotoxic potential of both T cell types was comparable. However, the killing pattern of ChoTCR-T cells was notably different from ChoTCR^CLU^-T cells. In contrast to ChoTCR^CLU^-T cells, ChoTCR-T cells destroyed tumor spheroids from the periphery, by forming large growing T cell clusters **(Figure 3C)**. This observation led us to hypothesize that ChoTCR^CLU^-T cells might have better infiltration capacity and kill more tumor cells in the spheroid core region. To test this, we designed another spheroid co-culture experiment using GFP^-^ A673 tumor cells and CellTrace CFSE-labelled T cells. This setup allowed us to visualize the distribution of T cells within the spheroid structure. The higher distribution of CFSE-positive ChoTCRCLU-T cells in the spheroid core following 3h co-culture implicated that OE of CLU might facilitate tumor infiltration capacity in these cells **(Figure 3D)**. This enhanced infiltration could potentially lead to more effective tumor cell killing throughout the spheroid. To further confirm this finding, we thoroughly washed out the tumor spheroid’s peripheral T cells and quantified spheroid-infiltrating T cells using flow cytometry **(Figure 3E)**. This method allowed us to specifically analyze the T cells that had penetrated the spheroid structure. The percentages of spheroid-infiltrating ChoTCR-positive cells were calculated as 3.71% and 6.29% for ChoTCR-T and ChoTCRCLU-T cells, respectively **(Figure 3F)**. This difference supports our hypothesis that CLU OE enhances T cell infiltration into tumor spheroids. As conventional microscopy only provides 2-dimensional spatial information about the distribution of T cells within the spheroid, we assessed A673 and TC71 spheroids by confocal microscopy. Conferring to our previous observations, we could verify that a higher number of ChoTCR^CLU^-T cells were able to persist within the spheroids. Interestingly, we observed a higher number of T cells undergoing apoptosis in the ChoTCR population than in NT and ChoTCR^CLU^-T cell populations **(Figure 3G-H)**. A 3-dimensional representation of z-stacked confocal images confirmed these findings **(Figure 3I)**. This data implies a major role of exhaustion and Activation-induced Cell death (AICD)^46^, both TCR-dependent programs that ultimately lead to apoptosis.

**Figure 3.**
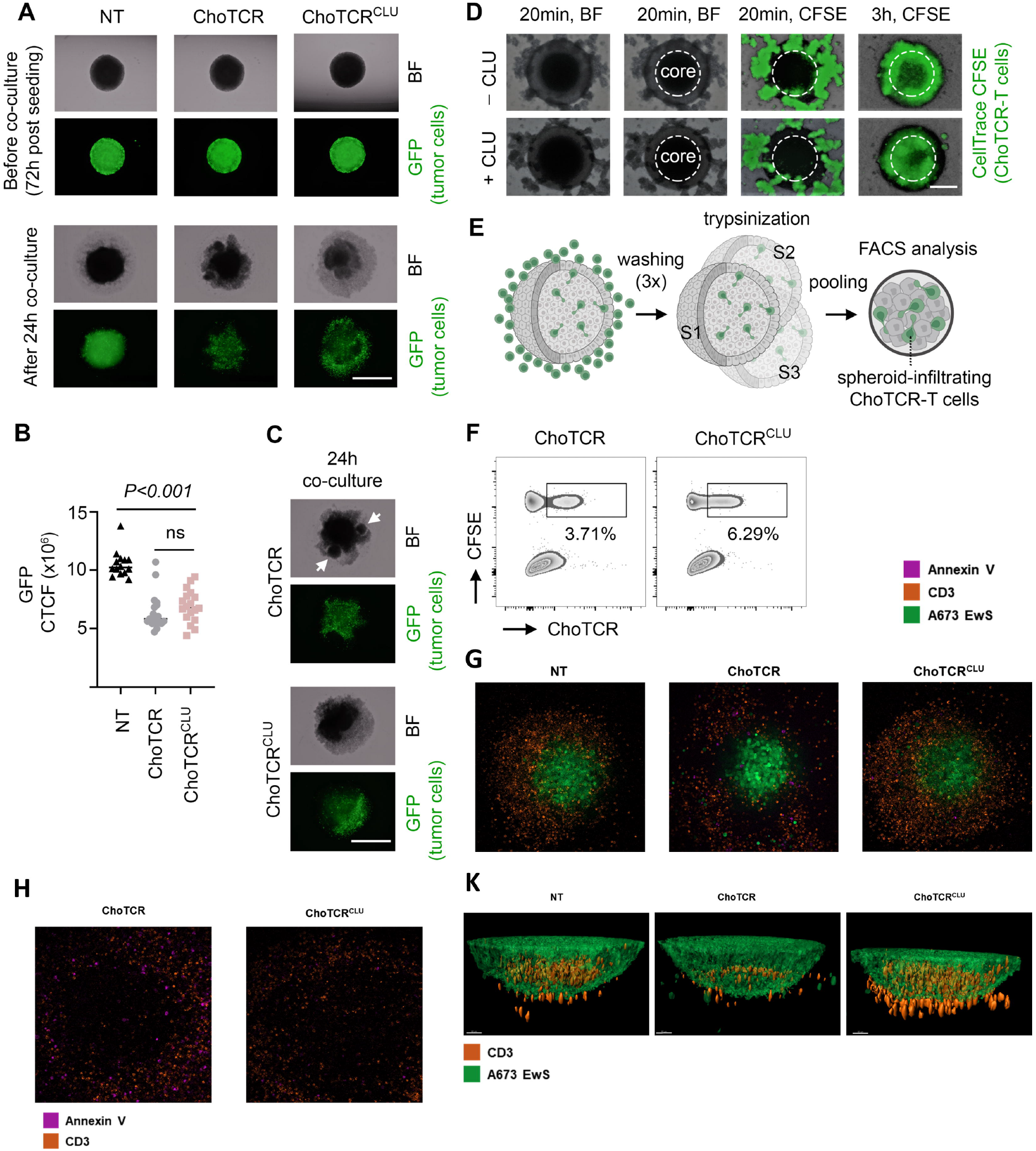
CLU OE facilitates tumor spheroid infiltration of ChoTCR-T cells. (**A**) Cytotoxicity effect of non-transduced (NT), ChoTCR-T, and ChoTCR^CLU^-T cells on the tumor spheroids generated from GFP-expressing A673 cells. Unlike NT cells, ChoTCR-T and ChoTCR^CLU^-T cells could destroy A673 tumor spheroids after 24h co-culture. BF: bright field. Scale bar=1000 µm. (**B**) Calculation of corrected total cell fluorescence (CTCF) based on the GFP intensity in the tumor spheroids co-cultured for 24h with NT (*n=14*), ChoTCR-T (*n=19*), and ChoTCR^CLU^-T cells (*n=18*). ns: non-significant. (**C**) Different pattern of spheroid destruction by ChoTCR-T and ChoTCR^CLU^-T cells after 24h co-culture. Arrows indicate the growing ChoTCR-T cell clusters in the spheroid’s peripheral zone. (**D**) Determination of spheroid infiltration capacity in the ChoTCR-T (-CLU) and ChoTCR^CLU^-T cells (+CLU) after 3h co-culture. Effector cells were labeled with CellTrace CFSE (green). The spheroid core is shown by dashed circles. Scale bar=400 µm. (**E**) Experiment design describing the preparation steps for flow cytometry analysis of A673 tumor spheroids after 5h co-culture with ChoTCR-T (*n=8*) or ChoTCR^CLU^-T cells (*n=9*). S: spheroid (**F**) Determination of spheroid-infiltrating ChoTCR-T or ChoTCR^CLU^-T cells by flow cytometry after 5h co-culture with A673 tumor spheroids. CellTrace CFSE was used to label effector cells, and CFSE+ChoTCR+ cells were detected within tumor spheroids. **(G)** Confocal microscopic image of A673 EwS spheroid infiltrating lymphocytes after 24hrs of co-culture at an E:T of 1:1. T cells were stained with a CD3 antibody before incubation of the co-culture (red), A673 cells were green, and AnnexinV surface expression was seen in purple. **(H)** Confocal microscopic image of TC71 EwS spheroid infiltrating lymphocytes after 24hrs of co-culture at an E:T of 1:1. CD3^+^ T cells are shown in red, AnnexinV surface expression in purple. **(I)** Three dimensional representations of z-stacked images acquired using a confocal microscope. A673 EwS Spheroids were imaged after 24 hours of co-culture at an E:T ratio of 1:1. CD3+ T cells are seen in red, and EwS cells are in green.

### 5.4 CLU OE Improves Persistence of T Cells *in vivo*

Our data so far has demonstrated reduced T cell exhaustion and increased persistence of ChoTCR^CLU^ T cells in tumor spheroids after prolonged incubation periods. In the next step, we evaluated the functional properties of CLU OE antigen-specific T cells in a tumor xenograft model. Luc-expressing A673 EwS cells were injected intramuscularly on Day 0, T cell treatments (NT, ChoTCR, and ChoTCR^CLU^) were administered on Day 8 **(Figure 4A)**. To evaluate the phenotype and persistence of the infused T cells, we analyzed engrafted human CD3^+^ cells in the spleen on Day 20 after injection of the tumor cells. In the ChoTCR group we found a significantly higher number of both AnnexinV^+^ and LAG3^+^cells, compared to the ChoTCR^CLU^ group, indicating an increased number of apoptotic cells and a faster transition of the ChoTCR-T cells into exhaustion. These observations indicate a markedly enhanced persistence of the ChoTCR^CLU^-T cells *in vivo*. Meanwhile, the NT control cells exhibited a much lower expression of these markers **(Figure 4B)**. In accordance with these results, we found an overall higher number of human CD3^+^-T cells in the spleens of the ChoTCR^CLU^-T cell treatment group. As the persistence of NT cells was comparable to the ChoTCR^CLU^-T cells, we assume the mechanism of the observed decline in T cell numbers to be, at least partly, dependent on TCR signaling **(Figure 4C)**. Likewise, there was an increased number of CD3^+^ cells in the mice’s lungs observed by immunohistochemistry in the CLU treatment group **(Figure 4D, BF**_10_**=1.41)**. *In vivo* imaging system (IVIS) measurements were performed on Days 7, 14, and 21 to evaluate the effect of different treatment approaches on tumor growth. Although it seemed like ChoTCR^CLU^-T cells where controlling tumor at later time points, no significant differences in tumor burden were observed between the treatment groups at any time point **(Figure 4E)**. To increase sensitivity, we compared the growth rates, defined as the fold change in signal intensity between measurements, during Week 1 and Week 2 post-infusion. Interestingly, ChoTCR^CLU^ showed a faster growth rate compared to ChoTCR during Week 1, while the growth rate was slower during Week 2. A Bayesian T-Test showed limited support for the alternative hypothesis (BF_10_=1.431) **(Figure 4F)**. Even though, we did not observe a prolonged survival of mice treated with tgT cells, immune histochemistry revealed a reduction of CD99^+^ single-cell EwS metastasis in lungs of mice treated with ChoTCR^CLU^-T cells **(Figure 4G-H)**. This preliminary evidence supports our *in vitro* observations.

**Figure 4.**
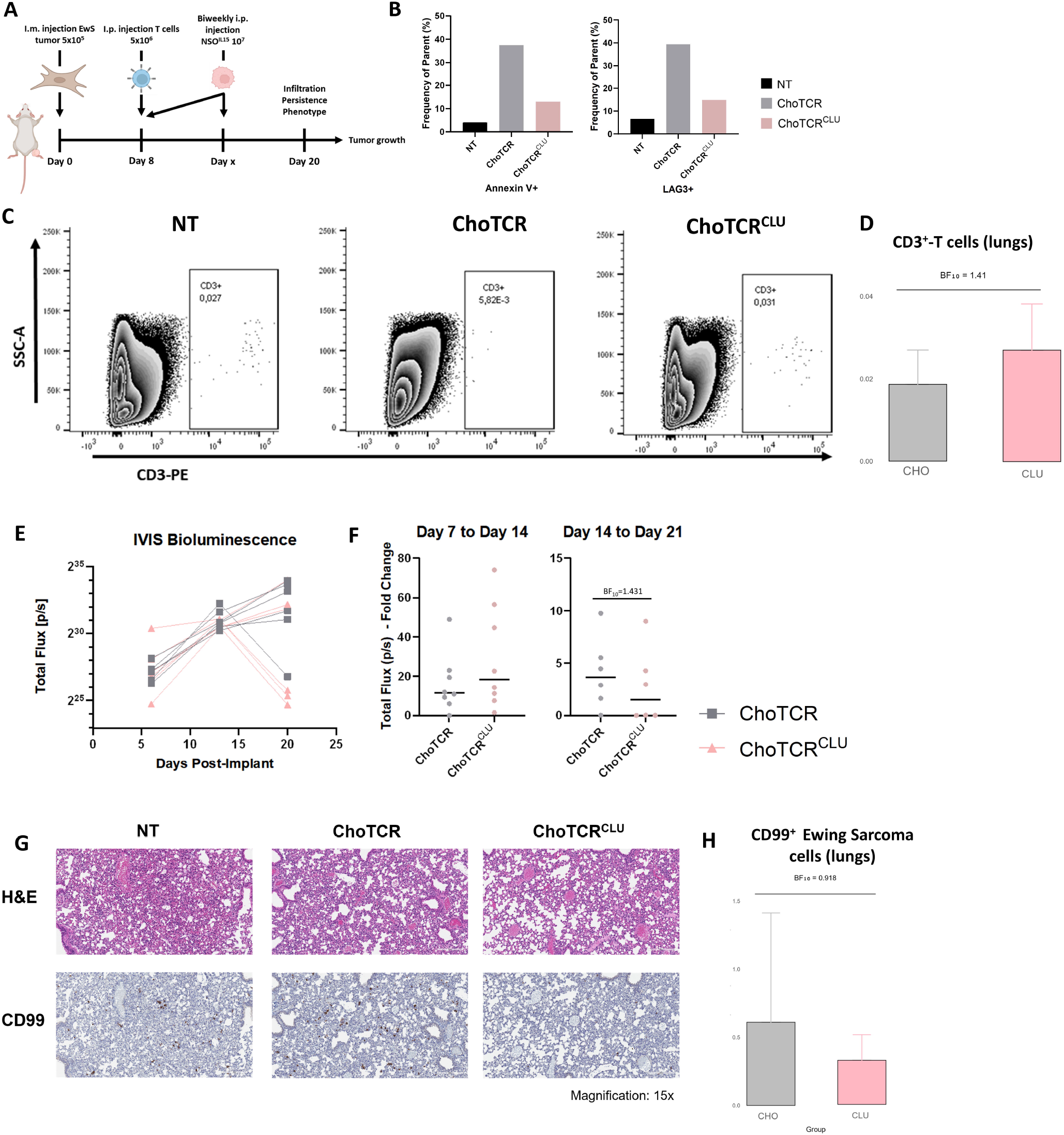
CLU induces improved persistence of ChoTCR-T cells *in vivo*. **(A)** Experimental Setup: 5×10^5^ A673 EwS cells were injected intramuscularly in the right calf of Rag2^−/−^ _γ_c^−/−^ mice at day 0. After 8 days, mice were randomized into treatment groups: NT, ChoTCR, and ChoTCR^CLU^. Mice were injected with 5×10^6^ T cells of the corresponding population and 10^7^ IL15-producing NSO cells previously irradiated at 100Gy. The application of NSO cells was repeated biweekly until the mice reached the predefined endpoint (defined by tumor diameter >1.7cm). **(B)** Expression of LAG3 and AnnexinV of T cells engrafted in the spleens of two representative animals at day 20. One representative blot for each condition is shown. **(C)** T cell persistence was analyzed in the spleens of two representative animals at day 20. One representative blot for each condition is shown. **(D)** CD3^+^ cells in lung tissue were quantified using QuPath. Positive cell detection was applied to identify CD3-expressing cells within the tissue sections.**(E)** Growth curves of EwS in mice measured by IVIS imaging. The graph shows bioluminescence flux (photons/second) of tumors in 6 individual mice at days 7, 14, and 21 after injection. Each line represents an individual mouse. **(F)** Comparison of growth rates measured by IVIS imaging depicted as fold change between Day 7 and Day 14, as well as between Day 14 and Day 21 after injection. **(G)** HE and CD99 IHC staining of mouse lungs at 15x magnification, shown for the NT, ChoTCR, and ChoTCR^CLU^ groups. Representative for three analyzed mouse lungs in each treatment group. **(H)** CD99^+^ cells in lung tissue were quantified using QuPath. Positive cell detection was applied to identify CD3-expressing cells within the tissue sections.

### 5.5 Proteomic Analysis Reveals Reduced Ribosomal Activity as a Possible Mechanism for Delayed Exhaustion

To elucidate the mechanism by which CLU OE delays T cell dysfunction, we compared NT, ChoTCR and ChoTCR^CLU^-T cells using TMT label-based quantitative mass spectrometry. Here, we observed 1388 differentially expressed proteins between ChoTCR and ChoTCR^CLU^, while 389 were upregulated and 999 were downregulated in ChoTCR^CLU^ **(Figure 5A)**. Among those proteins most significantly downregulated in ChoTCR^CLU^, we found gene products that were associated with ribosome biogenesis (e.g. RPL27, RPS3A, RPS9, RPS24, TRMT1) **(Figure 5B)**. Accordingly, the associated Gene ontology (GO) terms were enriched biological processes related to ribosomal RNA (rRNA) maturation, ribosome biogenesis and cytoplasmic translational activity **(Figure 5C)**. It has been shown that hyperactive protein synthesis, induced by a knock-out of the eucaryotic elongation factor 2 kinase (eEF2K) in T cells leads to increased energy consumption and thereby fuels the transition of T cells into a dysfunctional state and impairs their ability to infiltrate solid tumors.^9^ Accordingly, we saw a downregulation of the eucaryotic elongation factor 2 (eEF2), the canonical target of eEF2K, in ChoTCR^CLU^-T cells. We finally hypothesize, that CLU reduces global ribosome activity by stabilizing the posttranslational secondary structure of its targets, thereby reducing ER stress and altering cellular metabolism. Consistent with this hypothesis, we found a downregulation of several eukaryotic initiation factors (e.g. eIF5B, eIF3D, eIF2D) in ChoTCR^CLU^-T cells. These molecules are potent activators of translation by initiating the process of protein synthesis.^47^ Metabolism, in the context of T cell activation, strongly relies on sufficient glucose availability. Proliferating T cells rapidly switch from oxidative phosphorylation to aerobic glycolysis as their main source of energy.^48, 49^ Interestingly, in ChoTCR^CLU^ we observed a strong enrichment of GO terms associated with oligosaccharide and carbohydrate metabolism, which are important cellular suppliers of glucose **(Figure 5D-E)**.

**Figure 5.**
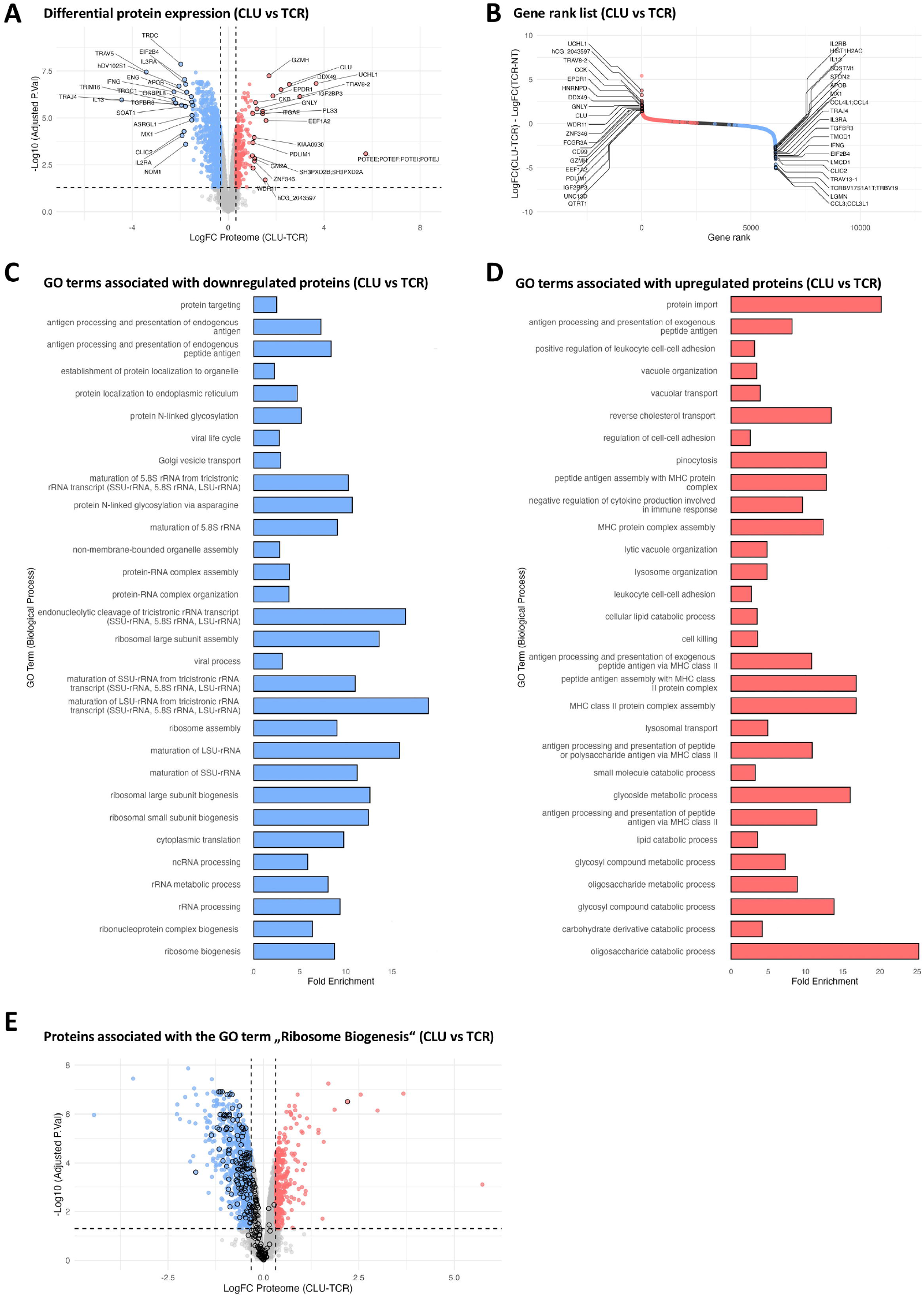
Proteomic analysis reveals reduced ribosome biogenesis and translation activity. **(A)** Volcano plot of proteins differentially regulated in ChoTCR vs. ChoTCR^CLU^-T cells. Proteins upregulated in ChoTCR^CLU^ are depicted as red, downregulated proteins are depicted as blue. **(B)** Log_2_ fold changes from **(A)** were plotted against their respected ranks. Highlighted are the top 20 proteins with the highest and lowest rank respectively. **(C) and (D)** Gene Ontology (GO) analysis of upregulated (red) and downregulated (blue) proteins in A, showing the top 30 significant pathways (FDR corrected p < 0.01). **(E)** Volcano plot of proteins differentially regulated in ChoTCR vs. ChoTCR^CLU^-T cells. Proteins associated with the GO term “Ribosome biogenesis” are highlighted with black circles.

### 5.6 Cytoprotective Gene Screening Identifies CLU as a Potential CAR-T cell enhancer

To identify cytoprotective genes potentially involved in safeguarding CAR-T cells from exhaustion, we reanalyzed publicly available bulk RNAseq expression data from Mediator of RNA polymerase II transcription, subunit 12 homolog (MED12) knockout (KO) GD2HAz CAR-T cells from the Stanford Mackall laboratory in a stimulated and unstimulated state. MED12 KO CAR-T cells exhibit robust resistance to T cell exhaustion and demonstrate increased efficacy *in vitro* and *in vivo*.^50, 51^ We screened 15 genes known to protect solid tumors from metabolic and therapy-induced stress. CLU was significantly overexpressed under both conditions **(Figure 6A-B)**. We also compared the expression of our candidate genes in a RNAseq dataset that included GD2 and CD19 CAR-T cells after 4h of stimulation with non-activated mock T cells. The majority of metabolic stress resistance genes in our screening were significantly upregulated, whereas CLU was markedly downregulated upon activation in both models **(Figure 6C-D)**. To assess post-transcriptional regulation of CLU in engineered T cells, we first co-cultured ChoTCR-T cells with A673 tumor cells and measured CLU expression via intracellular FACS staining 2, 4 and 6 days later. The expression of CLU in ChoTCR-T cells showed a significant decrease on day six. Likewise, CLU showed downregulation in GD2 and CD19 targeting CAR-T cells following five days of co-culture with GD2+ Nalm6 cells **(Figure 6E)**. Across different models,^50–53^ that have been established to improve CAR-T cell function, CLU was consistently upregulated, making it the standout candidate in this analysis **(Figure 6F)**. To assess the possible role of CLU in protecting CAR-T cells, we established CLU.GD2 CAR-T cells, using both 4-1BBζ and CD28ζ co-stimulatory domains. As GD2 CARs are tonically signalling, we ensured a comparable CAR expression in +/-CLU1 OE in both 4-1BBζ and CD28ζ bearing T cells, and similar cytotoxic activity when co-cultured with GD2^+^ NAlm6 tumor cells in short-term (24h and 48h) cocultures **(Figure 6G and Supplementary Figure 2C)**. Interestingly, the expression of inhibitory receptors CD39, and TIM3 was substantially higher in GD2 CAR-T cells compared to CLU.GD2 CAR-T cells **(Figure 6H and Supplementary Figure 2G)**. Conversely, when co-cultures were extended to 10 days, we observed a significant advantage of the CLU.GD2 CAR-T cells in both 4-1BBζ and CD28ζ models, as well as increased in IL-2 production **(Figure 6I-J)**. We also observed an increased functionality of CLU OE T cells under hypoxia, a potent trigger of T cell exhaustion **(Supplementary Figure 2D-F)**. This indicates that CLU OE may prevent the expression of inhibitory receptors, thereby indicating functional enhancement of CAR-T cells.

**Figure 6.**
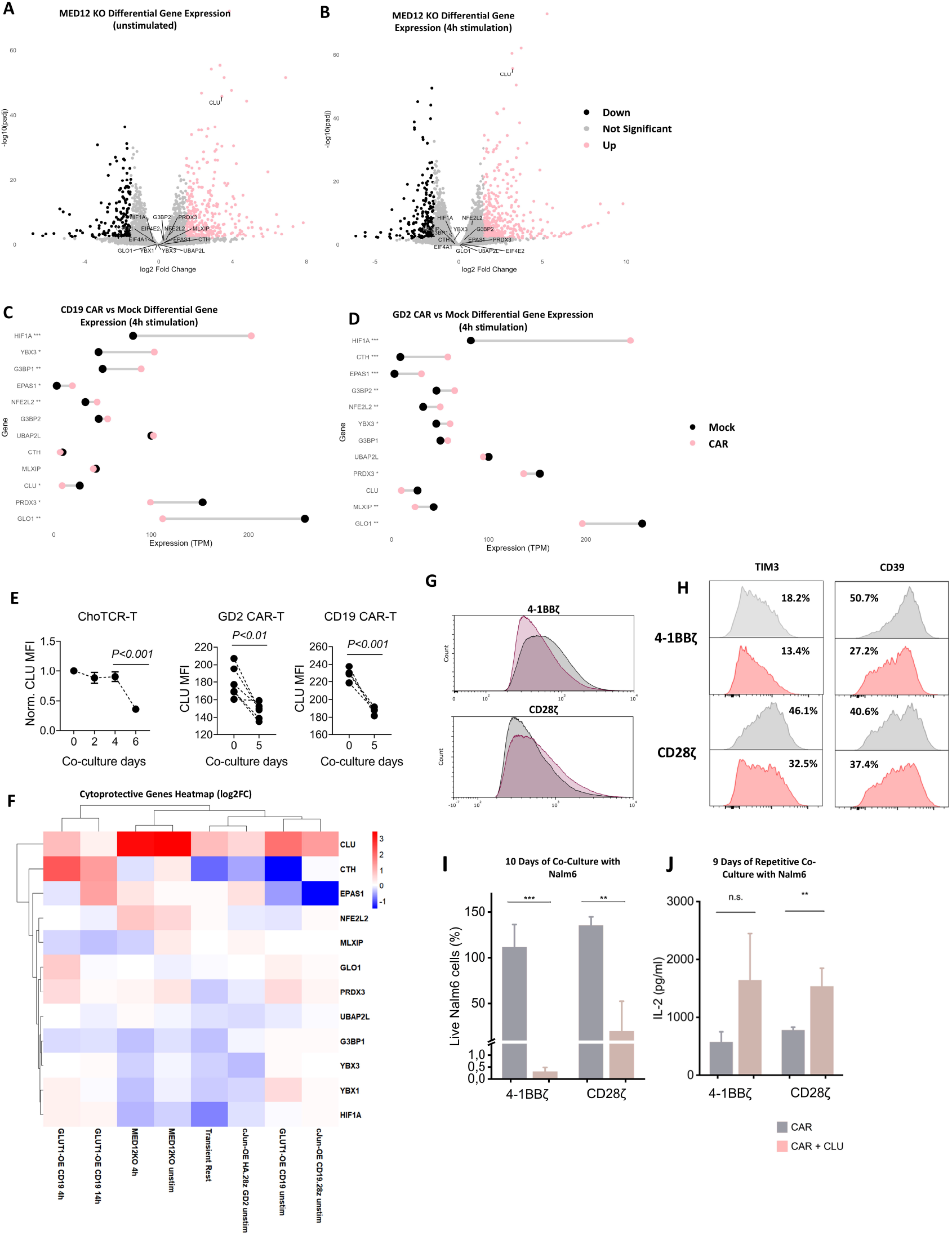
Expression of CLU in engineered T cells. (**A-B**) MED12 KO.GD2 CAR-T cells were stimulated for 4h/left unstimulated and then anlyzed by Bulk RNAseq. Cytoprotective genes as part of the screening were highlighted. Data reanalyzed from GSE174282 (**C-D)** Dumbbell plot representing the expression of screened cytoprotective genes in CD19.28z and HA.28z GD2 CAR-T cells after 4h stimulation compared to mock. Data reanalyzed after GSE275152 (**E**) CLU expression levels in the stimulated ChoTCR-T cells following co-culture with A673 tumor cells as indicated by normalized MFI (median fluorescence intensity). The MFI of CLU expression in the unstimulated ChoTCR-T cells was considered as control (0h). The effector-to-target ratio (E:T) is 1:10. CLU expression change in the CAR-T cells targeting GD2 (*n=6*) and CD19 (*n=4*) was assessed after five days of co-culture with target tumor cells. (**F**) Heatmap representing the log2FC expression change of the screened cytoprotective genes in established exhaustion-resistant CAR-T models. (**G**) Histogram representing the GD2-CAR expression on transduced human T cells after sorting for GFP expression. (**H**) Analysis of expression of CD39, and TIM3 in GD2 CAR-T cells after repetitive stimulation with GD2+ Nalm6 tumor cells for 9 days (E:T=1:10). (**I**) Luciferase (Luc) cytotoxicity assessment of GD-2 CAR-T cells after 10 days of co-culture with GD2^+^ Nalm6 cells (E:T 1:10). (**L**) Production of IL-2 measured via ELISA after repetitive stimulation with GD2+ Nalm6 tumor cells for 9 days (E:T=1:10).

## 6 Discussion

CAR-T cell therapies are a promising approach for treating hematologic malignancies and are increasingly being integrated into clinical practice. However, solid tumors present unresolved challenges due to their harsh tumor microenvironment (TME) and T cell exhaustion. To exploit the mechanisms tumors, use to adapt to the TME, we screened a single-cell RNA sequencing dataset for the expression of established cytoprotective genes across different T cell states. This analysis revealed a consistent downregulation of several such genes in exhausted T cells. Among them was MLXIP (MLX interacting protein), a gene known to contribute to the malignancy of pediatric B-lymphoblastic leukemia.^54^ In this study, we demonstrate that the heat-shock protein CLU is significantly downregulated upon long-term stimulation of tgTCR-T and CAR-T cells and generally in dysfunctional TILs. To stabilize and enhance CLU expression in T cells, we developed polycistronic constructs that induce transgenic expression of CLU in two distinct T cell models. While not impairing short-term functionality, OE of CLU led to a significant reduction in functional and phenotypic markers of T cell exhaustion following prolonged tumor cell stimulation. In 3D EwS spheroids and preliminary *in vivo* studies, ChoTCRCLU-T cells showed enhanced persistence and a trend toward reduced tumor growth. The observed high degree of variability might be explained by the relatively low number of mice in the experiment, as well as the already very efficacious Chm1 targeting TCR, which potentially overshadows the effect of the CLU OE. Mechanistically, TMT label-based quantitative mass spectrometry revealed that CLU OE alters the metabolic profile of T cells by downregulating ribosome biogenesis and translational activity. These findings align with prior evidence that translational activity is implicated in T cell exhaustion, as hyperactive translation increases energy consumption and promotes dysfunction.^9^ Our data indicate that CLU-mediated reductions in ribosome biogenesis involve the downregulation of key translation factors, including eEF2 and several eukaryotic initiation factors (eIFs), which are crucial for ribosome recruitment to mRNA. CLU reduces protein aggregation - a canonical role observed in cancer and neurodegenerative diseases - thereby alleviating ER stress and the demand for global protein synthesis.^14^ Assuming, that this cytoprotective effect in tumor cells is also relevant in T cells, we assume a CLU overexpressing T cell to have a slower turnover of proteins.

The modulation of T cell metabolism to enhance therapeutic durability is an attractive strategy in adoptive T cell therapies. OE of FOXO1^55^ and GLUT1^51^, as well as knock-out of eEF2K^9^, have been described to improve the functionality of adoptively transferred T cells by reprogramming their metabolism: either by reducing the cell’s need for nutrients or by increasing the availability of a nutrient to the cell.^17, 56^ Our findings suggest that CLU operates through a complementary mechanism, reducing the metabolic burden on T cells by stabilizing proteostasis and limiting excessive translational activity.

The TME of EwS has been described as an immune desert.^57, 58^ The number of TILs in the TME of EwS is a critical determinant of disease prognosis.^59^ The enhanced persistence and improved spheroid infiltration demonstrated in our study suggest potential therapeutic advantages of CLU-overexpressing T cells. Here, we observed a reduction of single-cell lung metastasis in mice after treatment with ChoTCRCLU-T cells. Given the results of the Bayesian estimate, we postulate that the prolonged functionality of CLU-overexpressing T cells might improve metastasis control. Considering, that the most significant cause of death in EwS (like in many other pediatric malignancies) is metastasis, targeting mechanisms that enhance metastasis control, could represent a promising therapeutic strategy to improve patient outcomes.^60^

In conclusion, our study highlights CLU OE as a promising strategy to address key limitations of CAR-T and tgTCR-T cell therapies in solid tumors. By potentially mitigating metabolic stress and exhaustion, CLU-overexpressing T cells demonstrate improved persistence and functionality, offering new avenues for enhancing the efficacy of adoptive T cell therapies. Further studies are warranted to validate these findings in larger-scale *in vivo* models and explore the implications of CLU-mediated metabolic reprogramming in engineered T cell platforms.

## 7 Author’s Contributions

CS: Formal analysis, Investigation, Visualization, Writing – original draft, Methodology, Data curation. YH: Investigation, Methodology. ES: Conceptualization, Writing – review & editing, Data curation, Formal analysis. AK: Investigation, Formal analysis, Methodology, Writing – review & editing. MP: Methodology, Investigation, Formal analysis. JH: Investigation, Methodology. ML: Investigation, Methodology. JM: Formal analysis, Visualization, Validation. FB: Formal analysis, Visualization, Validation. MS: Methodology, Writing – review & editing. BX: Methodology, Investigation. AM: Resources, Methodology. CB: Methodology, Investigation. AFM: Methodology, Investigation. JH: Resources. CM: Conceptualization, Writing – review & editing. JR: Conceptualization, Resources, Methodology, Supervision. PHS: Conceptualization, Resources, Supervision, Methodology. SEGB: Conceptualization, Project administration, Funding acquisition, Resources, Supervision, Writing – review & editing, Methodology.

## 8 Acknowledgements

The present work has been presented in part recently as a preliminary communication: Constantin Segner, Alisa Kolesnikova, Mansour Poorebrahim, Julia Höbart, Miriam Schulz, Busheng Xue, Christian Brückner, Alissia Fernandes Madeira, Julia Hauer, Jürgen Ruland, Poul H. Sorensen, Stefan EG Burdach. Clusterin overexpression in therapeutic T cell induces resistance to exhaustion [abstract]. In: Proceedings of the AACR Special Conference in Cancer Research: Advances in Pediatric Cancer Research; 2024 Sep 5-8; Toronto, Ontario, Canada. Philadelphia (PA): AACR; Cancer Res 2024;84(17 Suppl):Abstract nr B058.

## 9 Conflict of Interest Statement

S.E.G.B has ownership interest in PDL BioPharma and had US and EU intellectual properties in gene expression analysis. He served as consultant to EOS Biotechnology Inc. and serves as advisor to Bayer AG and Swedish Orphan Biovitrum AB. Other authors declare no conflict of interest.

## 10 Data Availability Statement

The mass spectrometry proteomics data have been deposited to the ProteomeXchange Consortium via the PRIDE partner repository with the dataset identifier PXD063762. The other data that support the findings of this study are available from the corresponding author, S.E.G.B, upon reasonable request.

## 11 Funding

The authors are grateful to the Cura Placida Children’s Cancer Research Foundation (CP101/22) for its financial support to CS, AK, CB and AFM. The Orbitrap Eclipse mass spectrometer was funded in part by the German Research Foundation (INST 95/1650-1 FUGG).

## 13 Tables

**Supplementary Table S1:**
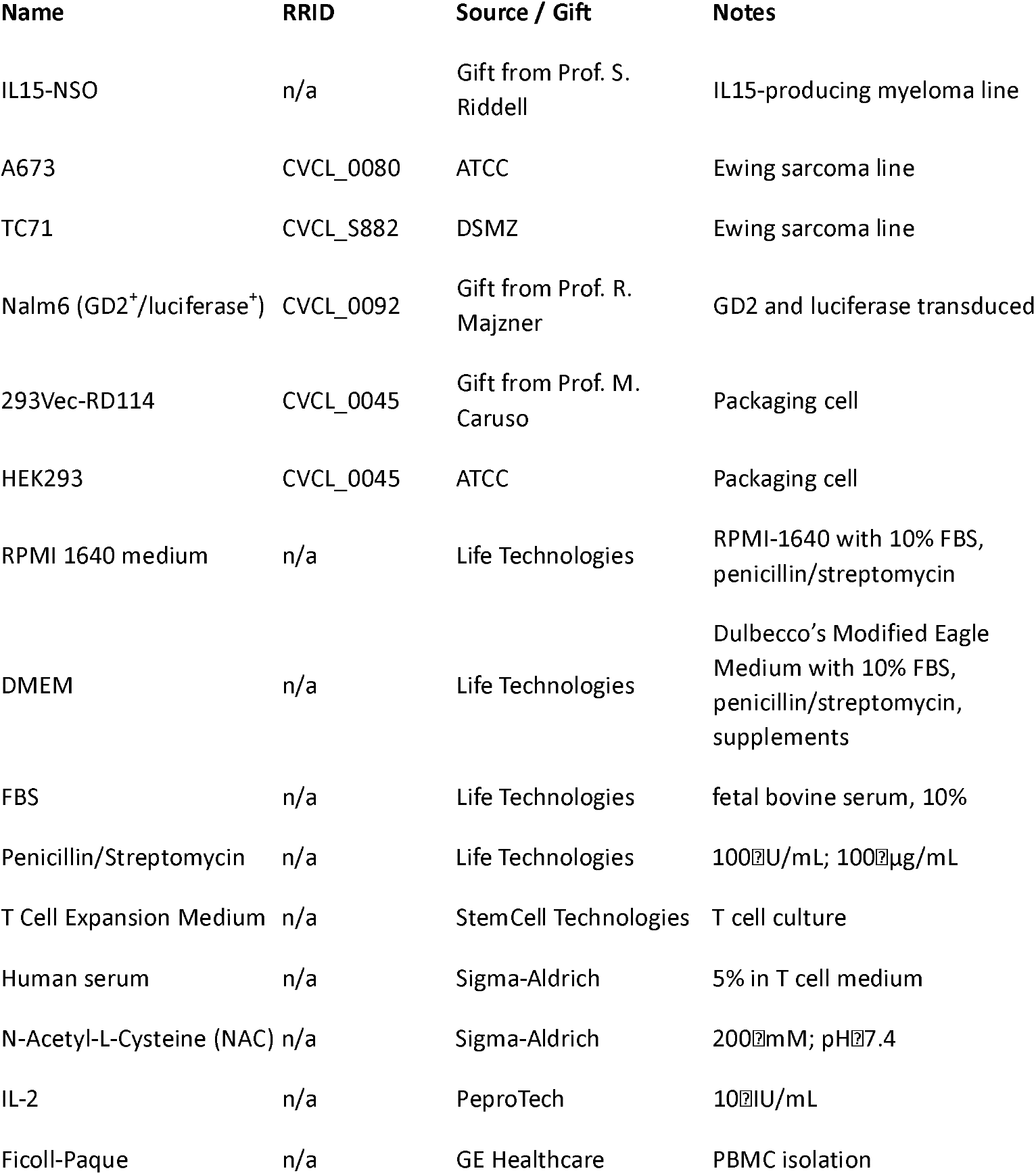
Cell lines, media and reagents.

**Supplementary Table S2:** Antibodies.

| **Name** | **Manufacturer** | **Clone** | **RRID** |
| --- | --- | --- | --- |
| Clusterin Antikörper (A-9): sc-166907 | Santa Cruz Biotechnology | A-9 (anti-Clusterin) | n.a. |
| PE/Cyanine5 anti-human CD279 (PD-1) Antibody | Biolegend | EH12.2H7 | AB_2910394 |
| FITC anti-human CD279 (PD-1) Antibody | Biolegend | EH12.2H7 | AB_940477 |
| Pacific Blue™ anti-human CD279 (PD-1) Antibody | Biolegend | EH12.2H7 | AB_2283437 |
| PE anti-human CD3 Antibody | Biolegend | HIT3a | AB_314044 |
| FITC anti-human CD3 Antibody | Biolegend | UCHT1 | AB_2562046 |
| Pacific Blue™ anti-human CD3 Antibody | Biolegend | UCHT1 | AB_2562048 |
| PerCP Cyanine 5.5 anti-human CD223 (LAG-3) Antibody | Biolegend | 11C3C65 | AB_2629755 |
| APC anti-human CD366 (Tim-3) Antibody | Biolegend | F38-2E2 | AB_2116574 |
| Alexa Fluor® 647 anti-human CD39 Antibody | Biolegend | A1 | n.a. |
| Alexa Fluor® 647 anti-human Annexin V Antibody | Biolegend | n.a. | n.a. |
| PE anti-mouse TCR β-chain | Biolegend | H57-597 | AB_313431 |
| Alexa Fluor® 647 anti-human Ki-67 | Biolegend | Ki-67 | AB_10900810 |
| Purified mouse anti-TOX | Biolegend | 6E6D03 | AB_2566643 |
| Purified rabbit anti-TCF1 (TCF7) | Biolegend | 7F11A10 | AB_2562103 |
| FITC anti-human/mouse Granzyme B Recombinant | Biolegend | QA16A02 | AB_2687029 |
| IgG Antibody, anti-human, FITC | Miltenyi Biotec | IS11-3B2.2.3 | AB_2733672 |
| IgG Antibody, anti-human, APC | Miltenyi Biotec | IS11-3B2.2.3 | AB_1036187 |
| IgG Antibody, anti-human, PE | Miltenyi Biotec | IS11-3B2.2.3 | n.a. |
| APC anti-human CD65L antibody | Miltenyi Biotec | 145/15 | AB_2733392 |

## 14 Figure Captions

**Supplementary Figure 1.**
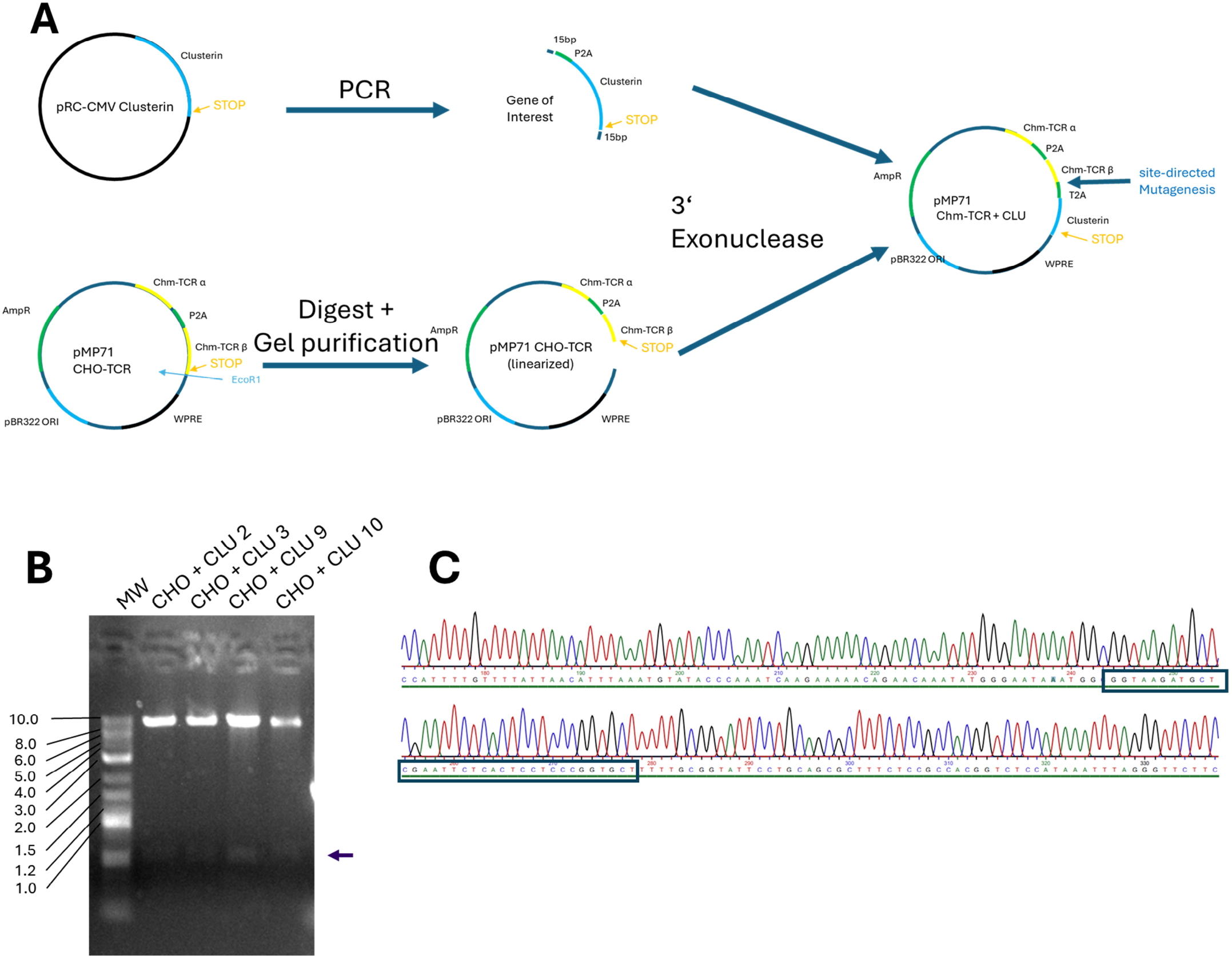
Cloning of the CLU gene into the pMP71 backbone: (**A**) Schematic representation of the cloning process using site-directed mutagenesis. (**B**) Gel electrophoresis showing the successfully transduced pMP71 ChoTCR + CLU plasmids after restriction digest with EcoRI. The second, smaller fragment of the plasmid is highlighted with an arrow (**C**) Chromatogram from the Sanger Sequencing of clone number 3 is highlighted with the black arrow. The insertion site of CLU at the 3’ end is highlighted with the blue box.

**Supplementary Figure 2:**
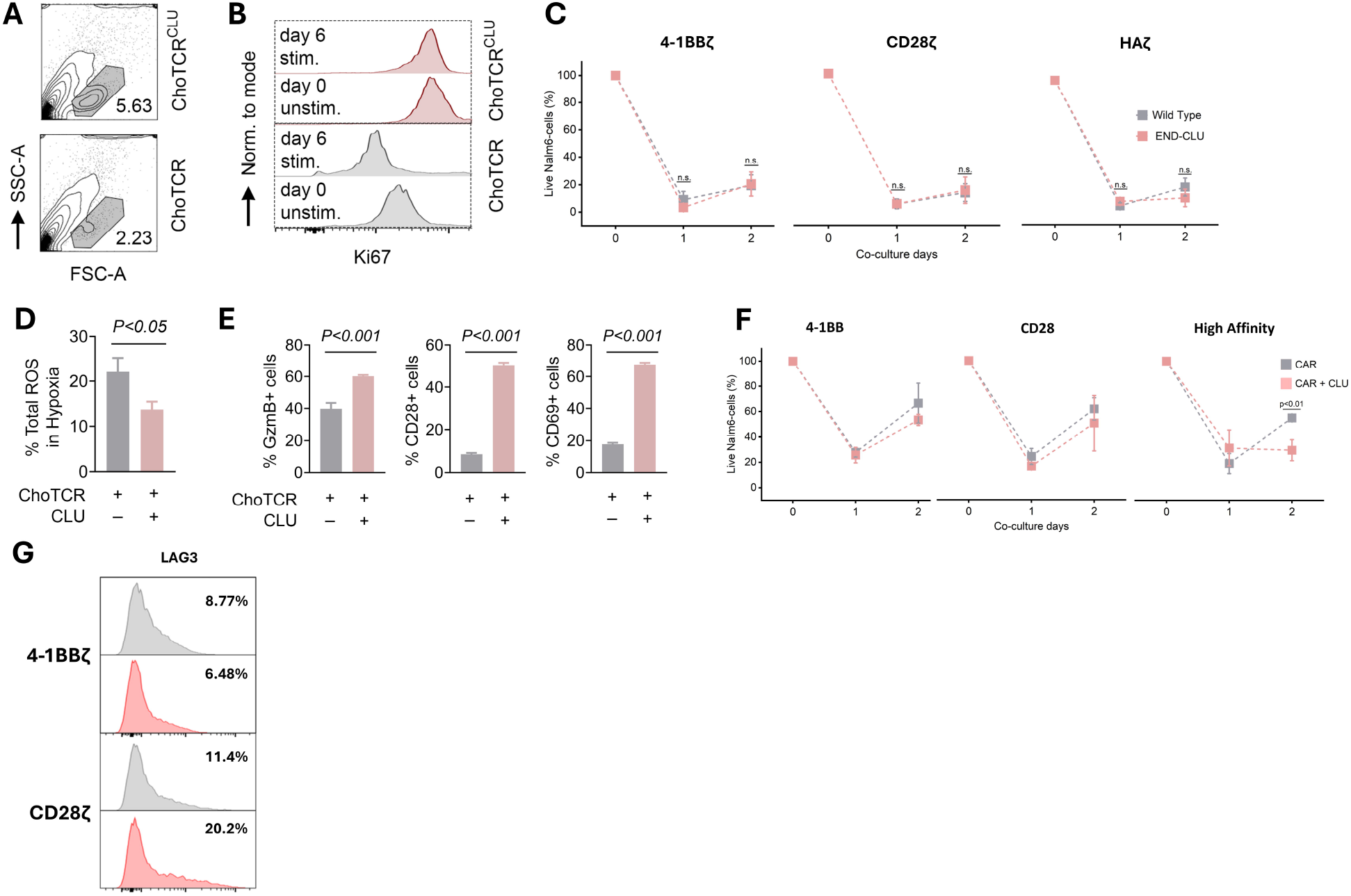
(**A**) Zebra Plot depicting lymphocyte size gate for ChoTCR-T cells after 6 days of co-culture with A673 EwS cells. (**B**) Histograms depicting Ki67 expression after 6 days co-culture with A673 cells. (**C**) Luciferase (Luc) cytotoxicity assessment of GD2 CAR-T cells after 24 and 48 hours of co-culture with GD2^+^ Nalm6 cells. (**D**)Assessment of ROS production in ChoTCR-T and ChoTCR^CLU^-T cells after 6h incubation under hypoxia (1.5% O_2_). (**E**) Expression of GzmB, CD28, and CD69 in ChoTCR-T and ChoTCRCLU-T cells after an overnight co-culture with A673 tumor cells in hypoxia condition (1.5% O2). In all experiments, T cells were firstly gated based on the ChoTCR expression. (**F**) Luciferase (Luc) cytotoxicity assessment of GD CAR-T cells after 24 and 48 hours of co-culture with GD2+ Nalm6 cells under hypoxia (1.5% O2). (**G**) Analysis of expression of LAG3 in GD2 CAR-T cells after repetitive stimulation with GD2+ Nalm6 tumor cells for 9 days (E:T=1:10).

## References

1. Hodi FS, O’Day SJ, McDermott DF, Weber RW, Sosman JA, Haanen JB, et al. Improved survival with ipilimumab in patients with metastatic melanoma. N Engl J Med. 2010;363(8):711–23.

2. Lipson EJ, Drake CG. Ipilimumab: an anti-CTLA-4 antibody for metastatic melanoma. Clin Cancer Res. 2011;17(22):6958–62.

3. Bourbon E, Ghesquieres H, Bachy E. CAR-T cells, from principle to clinical applications. Bull Cancer. 2021;108(10S):S4–S17.

4. Maude SL, Laetsch TW, Buechner J, Rives S, Boyer M, Bittencourt H, et al. Tisagenlecleucel in Children and Young Adults with B-Cell Lymphoblastic Leukemia. N Engl J Med. 2018;378(5):439–48.

5. Long AH, Haso WM, Shern JF, Wanhainen KM, Murgai M, Ingaramo M, et al. 4-1BB costimulation ameliorates T cell exhaustion induced by tonic signaling of chimeric antigen receptors. Nat Med. 2015;21(6):581–90.

6. Dolina JS, Van Braeckel-Budimir N, Thomas GD, Salek-Ardakani S. CD8(+) T Cell Exhaustion in Cancer. Front Immunol. 2021;12:715234.

7. Chow A, Perica K, Klebanoff CA, Wolchok JD. Clinical implications of T cell exhaustion for cancer immunotherapy. Nat Rev Clin Oncol. 2022;19(12):775–90.

8. Blank CU, Haining WN, Held W, Hogan PG, Kallies A, Lugli E, et al. Defining ‘T cell exhaustion’. Nat Rev Immunol. 2019;19(11):665–74.

9. Das JK, Ren Y, Kumar A, Peng HY, Wang L, Xiong X, et al. Elongation factor-2 kinase is a critical determinant of the fate and antitumor immunity of CD8(+) T cells. Sci Adv. 2022;8(5):eabl9783.

10. Wherry EJ. T cell exhaustion. Nat Immunol. 2011;12(6):492–9.

11. Hashimoto M, Kamphorst AO, Im SJ, Kissick HT, Pillai RN, Ramalingam SS, et al. CD8 T Cell Exhaustion in Chronic Infection and Cancer: Opportunities for Interventions. Annu Rev Med. 2018;69:301–18.

12. Ruffo E, Wu RC, Bruno TC, Workman CJ, Vignali DAA. Lymphocyte-activation gene 3 (LAG3): The next immune checkpoint receptor. Semin Immunol. 2019;42:101305.

13. Xie X, Feng Y, Fan P, Dong D, Yao X, Peng Y, et al. Increased co-expression of TIM-3 with TIGIT or 2B4 on CD8+ T cells is associated with poor prognosis in locally advanced nasopharyngeal carcinoma. Biomol Biomed. 2023;23(4):584–95.

14. Rodriguez-Rivera C, Garcia MM, Molina-Alvarez M, Gonzalez-Martin C, Goicoechea C. Clusterin: Always protecting. Synthesis, function and potential issues. Biomed Pharmacother. 2021;134:111174.

15. Wiggs JL, Kang JH, Fan B, Levkovitch-Verbin H, Pasquale LR. A Role for Clusterin in Exfoliation Syndrome and Exfoliation Glaucoma? J Glaucoma. 2018;27 Suppl 1(Suppl 1):S61–S6.

16. Carver JA, Rekas A, Thorn DC, Wilson MR. Small heat-shock proteins and clusterin: intra- and extracellular molecular chaperones with a common mechanism of action and function? IUBMB Life. 2003;55(12):661–8.

17. Garcia-Aranda M, Tellez T, Munoz M, Redondo M. Clusterin inhibition mediates sensitivity to chemotherapy and radiotherapy in human cancer. Anticancer Drugs. 2017;28(7):702–16.

18. Scharping NE, Rivadeneira DB, Menk AV, Vignali PDA, Ford BR, Rittenhouse NL, et al. Mitochondrial stress induced by continuous stimulation under hypoxia rapidly drives T cell exhaustion. Nat Immunol. 2021;22(2):205–15.

19. Huang Y, Si X, Shao M, Teng X, Xiao G, Huang H. Rewiring mitochondrial metabolism to counteract exhaustion of CAR-T cells. J Hematol Oncol. 2022;15(1):38.

20. Spector LG, Hubbard AK, Diessner BJ, Machiela MJ, Webber BR, Schiffman JD. Comparative international incidence of Ewing sarcoma 1988 to 2012. Int J Cancer. 2021;149(5):1054–66.

21. Jawad MU, Cheung MC, Min ES, Schneiderbauer MM, Koniaris LG, Scully SP. Ewing sarcoma demonstrates racial disparities in incidence-related and sex-related differences in outcome: an analysis of 1631 cases from the SEER database, 1973-2005. Cancer. 2009;115(15):3526–36.

22. Jardim DL, Goodman A, de Melo Gagliato D, Kurzrock R. The Challenges of Tumor Mutational Burden as an Immunotherapy Biomarker. Cancer Cell. 2021;39(2):154–73.

23. Thiel U, Schober SJ, Einspieler I, Kirschner A, Thiede M, Schirmer D, et al. Ewing sarcoma partial regression without GvHD by chondromodulin-I/HLA-A<sup>*</sup>02:01-specific allorestricted T cell receptor transgenic T cells. Oncoimmunology. 2017;6(5):e1312239.

24. Staege MS, Hutter C, Neumann I, Foja S, Hattenhorst UE, Hansen G, et al. DNA microarrays reveal relationship of Ewing family tumors to both endothelial and fetal neural crest-derived cells and define novel targets. Cancer Res. 2004;64(22):8213–21.

25. Hiraki Y, Inoue H, Iyama K, Kamizono A, Ochiai M, Shukunami C, et al. Identification of chondromodulin I as a novel endothelial cell growth inhibitor. Purification and its localization in the avascular zone of epiphyseal cartilage. J Biol Chem. 1997;272(51):32419–26.

26. von Heyking K, Calzada-Wack J, Gollner S, Neff F, Schmidt O, Hensel T, et al. The endochondral bone protein CHM1 sustains an undifferentiated, invasive phenotype, promoting lung metastasis in Ewing sarcoma. Mol Oncol. 2017;11(9):1288–301.

27. van der Schaft DW, Hillen F, Pauwels P, Kirschmann DA, Castermans K, Egbrink MG, et al. Tumor cell plasticity in Ewing sarcoma, an alternative circulatory system stimulated by hypoxia. Cancer Res. 2005;65(24):11520–8.

28. Ewing J. Classics in oncology. Diffuse endothelioma of bone. James Ewing. Proceedings of the New York Pathological Society, 1921. CA Cancer J Clin. 1972;22(2):95–8.

29. Nazha B, Inal C, Owonikoko TK. Disialoganglioside GD2 Expression in Solid Tumors and Role as a Target for Cancer Therapy. Front Oncol. 2020;10:1000.

30. Spasov NJ, Dombrowski F, Lode HN, Spasova M, Ivanova L, Mumdjiev I, et al. First-line Anti-GD2 Therapy Combined With Consolidation Chemotherapy in 3 Patients With Newly Diagnosed Metastatic Ewing Sarcoma or Ewing-like Sarcoma. J Pediatr Hematol Oncol. 2022;44(6):e948–e53.

31. Blaeschke F, Thiel U, Kirschner A, Thiede M, Rubio RA, Schirmer D, et al. Human HLA-A<sup>*</sup>02:01/CHM1+ allo-restricted T cell receptor transgenic CD8+ T cells specifically inhibit Ewing sarcoma growth in vitro and in vivo. Oncotarget. 2016;7(28):43267–80.

32. Xue B, von Heyking K, Gassmann H, Poorebrahim M, Thiede M, Schober K, et al. T Cells Directed against the Metastatic Driver Chondromodulin-1 in Ewing Sarcoma: Comparative Engineering with CRISPR/Cas9 vs. Retroviral Gene Transfer for Adoptive Transfer. Cancers (Basel). 2022;14(22).

33. Bankhead P. QuPath: Open source software for digital pathology image analysis. Version 0.5.1 ed: University of Edinburgh; 2017.

34. Ruprecht B, Zecha J, Zolg DP, Kuster B. High pH Reversed-Phase Micro-Columns for Simple, Sensitive, and Efficient Fractionation of Proteome and (TMT labeled) Phosphoproteome Digests. Methods Mol Biol. 2017;1550:83–98.

35. Tyanova S, Temu T, Cox J. The MaxQuant computational platform for mass spectrometry-based shotgun proteomics. Nat Protoc. 2016;11(12):2301–19.

36. Bryan HWaJ. readxl: Read Excel Files. 2023.

37. Wickham H. ggplot2: Elegant Graphics for Data Analysis: Springer-Verlag New York; 2016.

38. Vaughan HWaRFaLHaKMaD. dplyr: A Grammar of Data Manipulation. 2023.

39. Slowikowski K. ggrepel: Automatically Position Non-Overlapping Text Labels with ‘ggplot2’. 2024.

40. Monaco G, Lee B, Xu W, Mustafah S, Hwang YY, Carré C, et al. RNA-Seq Signatures Normalized by mRNA Abundance Allow Absolute Deconvolution of Human Immune Cell Types. Cell Rep. 2019;26(6):1627-40.e7.

41. FlowJo. 10.9 ed: BD Life Sciences; 2024.

42. OriginPro 2024. 2024 ed: OriginLab Corporation; 2024.

43. GraphPad Prism. 10.0 ed: GraphPad Software; 2023.

44. Wittwer J, Bradley D. Clusterin and Its Role in Insulin Resistance and the Cardiometabolic Syndrome. Front Immunol. 2021;12:612496.

45. Park S, Mathis KW, Lee IK. The physiological roles of apolipoprotein J/clusterin in metabolic and cardiovascular diseases. Rev Endocr Metab Disord. 2014;15(1):45–53.

46. Richter GH, Mollweide A, Hanewinkel K, Zobywalski C, Burdach S. CD25 blockade protects T cells from activation-induced cell death (AICD) via maintenance of TOSO expression. Scand J Immunol. 2009;70(3):206–15.

47. Sonenberg N, Hinnebusch AG. Regulation of translation initiation in eukaryotes: mechanisms and biological targets. Cell. 2009;136(4):731–45.

48. Wang T, Marquardt C, Foker J. Aerobic glycolysis during lymphocyte proliferation. Nature. 1976;261(5562):702–5.

49. Greiner EF, Guppy M, Brand K. Glucose is essential for proliferation and the glycolytic enzyme induction that provokes a transition to glycolytic energy production. J Biol Chem. 1994;269(50):31484–90.

50. Freitas KA, Belk JA, Sotillo E, Quinn PJ, Ramello MC, Malipatlolla M, et al. Enhanced T cell effector activity by targeting the Mediator kinase module. Science. 2022;378(6620):eabn5647.

51. Guerrero JA, Klysz DD, Chen Y, Malipatlolla M, Lone J, Fowler C, et al. GLUT1 overexpression in CAR-T cells induces metabolic reprogramming and enhances potency. Nat Commun. 2024;15(1):8658.

52. Weber EW, Parker KR, Sotillo E, Lynn RC, Anbunathan H, Lattin J, et al. Transient rest restores functionality in exhausted CAR-T cells through epigenetic remodeling. Science. 2021;372(6537).

53. Lynn RC, Weber EW, Sotillo E, Gennert D, Xu P, Good Z, et al. c-Jun overexpression in CAR T cells induces exhaustion resistance. Nature. 2019;576(7786):293–300.

54. Sipol A, Hameister E, Xue B, Hofstetter J, Barenboim M, Ollinger R, et al. MondoA drives malignancy in B-ALL through enhanced adaptation to metabolic stress. Blood. 2022;139(8):1184–97.

55. Chan JD, Scheffler CM, Munoz I, Sek K, Lee JN, Huang YK, et al. FOXO1 enhances CAR T cell stemness, metabolic fitness and efficacy. Nature. 2024;629(8010):201–10.

56. Gregory JM, Whiten DR, Brown RA, Barros TP, Kumita JR, Yerbury JJ, et al. Clusterin protects neurons against intracellular proteotoxicity. Acta Neuropathol Commun. 2017;5(1):81.

57. Brohl AS, Sindiri S, Wei JS, Milewski D, Chou HC, Song YK, et al. Immuno-transcriptomic profiling of extracranial pediatric solid malignancies. Cell Rep. 2021;37(8):110047.

58. Evdokimova V, Gassmann H, Radvanyi L, Burdach SEG. Current State of Immunotherapy and Mechanisms of Immune Evasion in Ewing Sarcoma and Osteosarcoma. Cancers (Basel). 2022;15(1).

59. Cillo AR, Mukherjee E, Bailey NG, Onkar S, Daley J, Salgado C, et al. Ewing Sarcoma and Osteosarcoma Have Distinct Immune Signatures and Intercellular Communication Networks. Clin Cancer Res. 2022;28(22):4968–82.

60. Grunewald TGP, Cidre-Aranaz F, Surdez D, Tomazou EM, de Alava E, Kovar H, et al. Ewing sarcoma. Nat Rev Dis Primers. 2018;4(1):5.

